# Fine-Tuning *TAWAWA1*-Mediated Panicle Architecture by Genome Editing of a Downstream Conserved Noncoding Sequence in Rice

**DOI:** 10.1101/2024.09.17.612847

**Authors:** Takeshi Kuroha, Fabien Lombardo, Watal M Iwasaki, Svetlana Chechetka, Yoshihiro Kawahara, Akiko Yoshida, Junko Kyozuka, Takashi Makino, Hitoshi Yoshida

## Abstract

Genome editing is a promising tool to enhance plant breeding, particularly for the generation of practical quantitative traits. We focused on the rice yield-related gene *TAWAWA1* (*TAW1*) and produced various degrees of increased panicle branching phenotypes by modifying its downstream conserved noncoding sequence (CNS). Differential modification of two highly conserved regions containing putative repressive elements in this CNS caused transcriptional upregulation of *TAW1*, resulting in diverse phenotypes.

## Main text

Genome editing has significantly advanced in recent years, with numerous attempts to integrate these technologies into crop breeding ^1^. Many useful agronomic traits result from subtle changes in gene expression patterns conferred by natural variations ^2^. Therefore, the modification of regulatory sequences through genome editing presents a potential strategy to develop practical breeding resources. Promoters and/or *cis*-regulatory elements of several target genes have been extensively edited to alter their gene expression patterns in tomato ^3^, maize ^4^, and rice ^5–7^. However, this approach requires numerous genome edits across a wide range of promoter regions. The selection of optimal target sites for genome editing from large intragenic noncoding regions to alter gene expression patterns and produce ideal agronomic traits remains challenging. Here, we report the production of quantitative trait variations in panicle branching through precise genome editing of a conserved noncoding sequence (CNS) located downstream of the rice yield-related gene *TAWAWA1* (*TAW1*) ^8^. Different patterns of CNS modification caused transcriptional upregulation of *TAW1*, which led to various degrees of increased panicle branching phenotypes.

*TAW1* is a member of the ALOG (*Arabidopsis* LSH1 and *Oryza* G1) gene family encoding putative transcriptional regulators. In grass species, ALOG family proteins are essential for specification of floral organ identity and the normal development of spikelet and inflorescence architecture ^8–10^. In a screen of a transposon-mutagenized rice population, Yoshida *et al*. ^8^ isolated two allelic mutants, *taw1*-*D1* and *taw1*-*D2*, which exhibited elevated *TAW1* expression and increased panicle branching. Both mutant lines carried *nDart1*-*0* transposons inserted approximately 700 bp downstream from the transcriptional termination site of *TAW1* (Fig. 1a) (ref. 8). Although the insertion sites differed by only 16 bp in the respective mutants, the inflorescences of *taw1*-*D1* were considerably more branched than those of *taw1*-*D2* (ref. 8), suggesting that the precise insertion site determines the branching phenotype severity. Because genes responsible for inflorescence architecture are largely conserved in grass species, we hypothesized that conserved regulatory sequences might exist around the *taw1-D1/-D2* insertion sites in these species. We first identified *TAW1* homologues in monocot species through phylogenetic analysis (Supplementary Fig. 1a, b), and then compared their genomic sequences using mVISTA to detect conserved sequences (Fig. 1a). We subsequently identified a CNS (hereafter, *TAW1*-CNS) in grass species, including the BEP clade, located 32 and 48 bp downstream of the transposon insertion sites in *taw1-D1* and *taw1-D2*, respectively (Fig. 1b and Supplementary Fig. 2). RNA sequencing (RNA-seq) analysis of young panicles did not detect transcription from the *TAW1*-CNS region, suggesting that *TAW1* upregulation in the transposon alleles was not caused by antisense-related mechanisms (Supplementary Fig. 3).

**Fig. 1.**
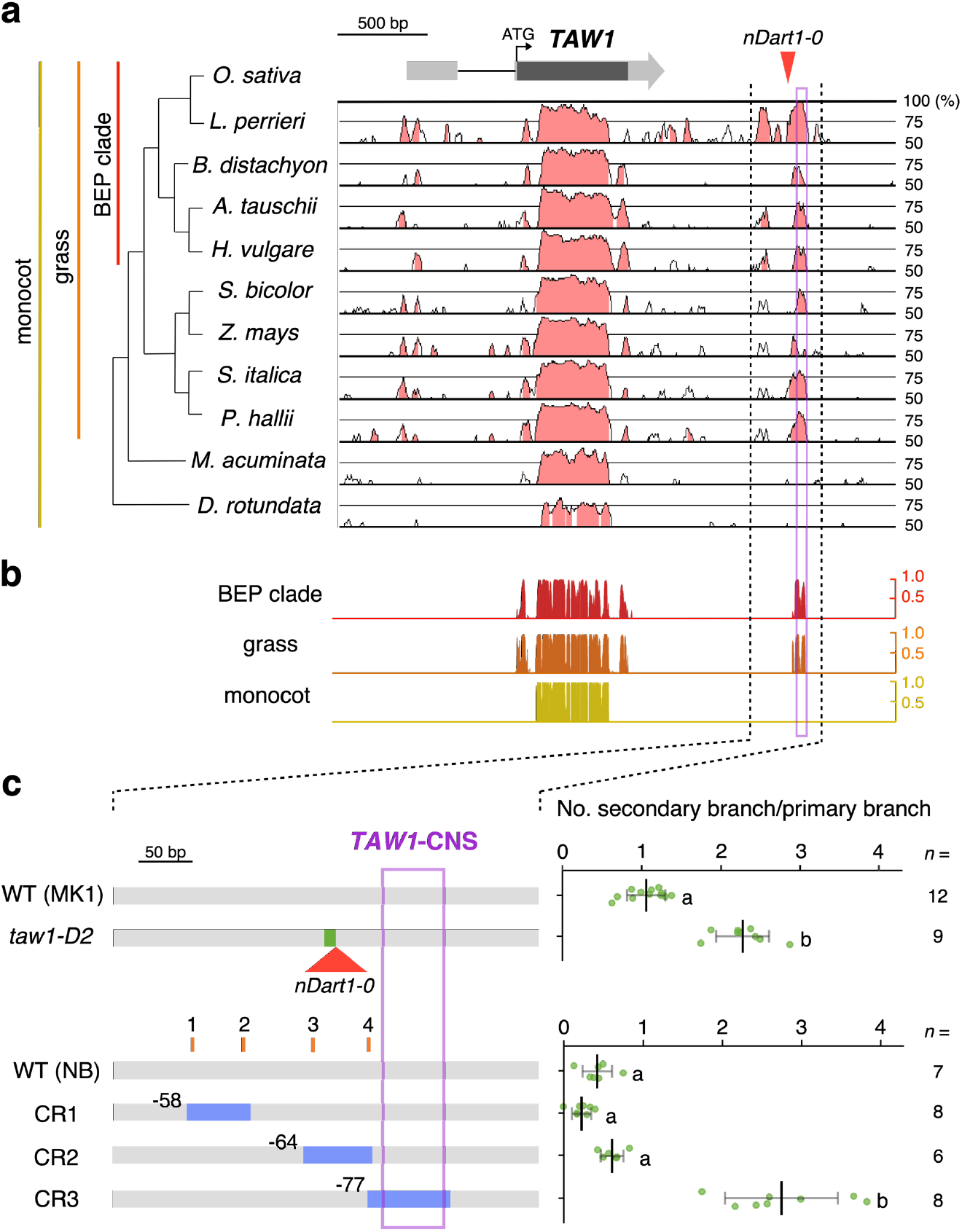
Deletion of the *TAW1*-CNS region increases panicle branching in rice. **a**, mVISTA plots of the *TAW1* genomic region. The rice *TAW1* gene structure is shown at the top. The grey box, grey arrow, black box, and horizontal line represent the 5′-UTR, 3′-UTR, exon, and intron of the *TAW1* gene, respectively. mVISTA plots of genomic region sequences across *TAW1* of rice and *TAW1* homologues from other monocot species are shown. Regions of high similarity defined by mVISTA are highlighted in pink. **b**, Similarity of the *TAW1* genomic region within monocot, grass, and BEP clade groups. **c**, Panicle branching phenotypes of *taw1-D2* (upper panel) and genome-edited lines (lower panel). The rice genomic region between the two dotted lines in **a** and **b** is magnified in the left panels. Positions of duplications and insertions in *taw1-D2* are indicated in the upper left panel as a green square and a red triangle, respectively. Positions of deletions introduced by genome editing are indicated in the lower left panel as blue boxes. PAM sequence positions in gRNA targets for genome editing (1 to 4) are shown as orange lines. MK1 and Nipponbare (NB) were used as WT for the *taw1-D* and genome-edited lines, respectively. Different letters in the right panels indicate statistically significant differences (*P* < 0.05; Tukey–Kramer test). *TAW1*-CNS region detected in the BEP clade is indicated as a purple box in **a**–**c**.

To elucidate the function of *TAW1*-CNS, we produced genome-edited lines with deletions of *TAW1* downstream regions using the clustered regularly interspaced short palindromic repeats (CRISPR)–CRISPR-associated protein-9 nuclease (Cas9) system and compared their phenotypes with the *taw1-D2* mutant. Whereas deletion of the genomic region including *taw1-D1/-D2* insertion sites or regions further upstream resulted in a normal phenotype, deletion of the genomic region encompassing the entire *TAW1*-CNS significantly increased the number of secondary branches per primary branch (i.e., panicle branching) (Fig. 1c). This increased branching phenotype was similar to the *taw1-D2* mutant (Fig. 1c), strongly suggesting that the phenotypes of *taw1-D1/-D2* alleles are not caused by potential enhancer-like activity of the *nDart1-0* transposon, but by transposon insertion-mediated disturbance of transcriptional repression activity of the *TAW1*-CNS region itself. The observation that the transposon insertion site in *taw1-D1* was closer to *TAW1*-CNS, relative to the insertion site in *taw1-D2* (Supplementary Fig. 2) (ref. 8), suggested that more proximal insertion has a stronger effect in disturbing the putative repression activity of *TAW1*-CNS. Semi-dominant inheritance of the panicle branching phenotype in the *TAW1*-CNS deletion mutant (Supplementary Fig. 4), consistent with the inheritance of *taw1-D1/-D2* traits ^8^, further supported the hypothesis of the repressive function of *TAW1*-CNS. We next generated additional lines with genome edits around *TAW1*-CNS (Fig. 2a–c) and compared transcription levels of *TAW1* and *TAW1*-proximal genes in immature inflorescence of genome-edited lines with moderate (CR19) and severe (CR3) phenotypes by RNA-seq (Fig. 2d). Whereas *TAW1* transcription levels were positively correlated with panicle branching phenotype strength (Fig. 2b–d), the genome-edited lines showed no significant differences in transcription levels of *TAW1*-proximal genes (Fig. 2d, Supplementary Fig. 5). This indicates that *TAW1*-CNS acts as a gene-specific silencer of *TAW1* in the wild-type (WT) background.

**Fig. 2.**
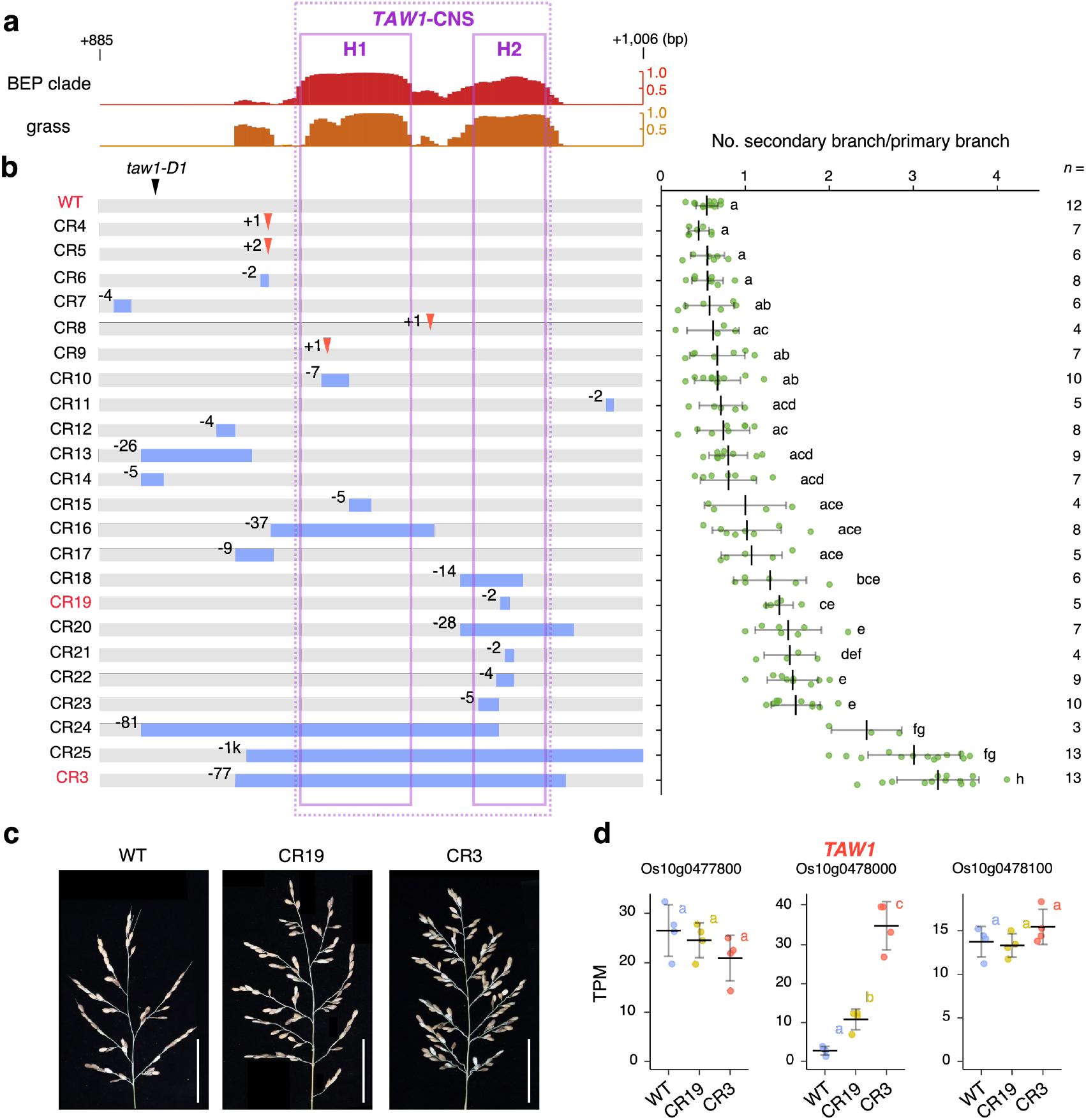
Creation of quantitative trait variations with transcriptional upregulation of *TAW1* by precise genome editing of *TAW1*-CNS. **a**, Similarity of the *TAW1*-CNS region in grass or BEP clade groups. The region 885–1006 bp downstream from the *TAW1* stop codon in *O. sativa* is shown. **b**, Panicle branching phenotypes of genome-edited lines. Positions of deletions and insertions are indicated in the left panel as blue boxes and red triangles, respectively. Different letters in the right panel indicate statistically significant differences (*P* < 0.05; Tukey–Kramer test). **c**, Panicle morphology of WT (Nipponbare), CR19, and CR3 genome-edited lines. Scale bars: 5 cm. **d**, Normalized transcript expression levels (TPM) of *TAW1* and proximal genes in immature inflorescence detected by RNA-seq analysis. Different letters indicate statistically significant differences (*P* < 0.05; Tukey–Kramer test) (*n* = 4). Detailed results are provided in Supplementary Fig. 5. Dashed box in the left panel indicates *TAW1*-CNS in **a** and **b**. Solid squares indicate highly conserved CNS regions with conservation score ≥ 0.6 in the BEP clade in **a** and **b**.

We identified two highly conserved regions (H1 and H2) in *TAW1*-CNS, with conservation scores ≥ 0.6 in the BEP clade (Supplementary Fig. 6). To precisely evaluate the functions of *TAW1*-CNS, including H1 and H2, in panicle branching, we compared the phenotypes of genome-edited lines with different mutations in, across, or proximal to the CNS, harbouring small indels or large deletions (Fig. 2a, b, and Supplementary Fig. 7). We confirmed that genome-edited lines with large deletions of all or substantial parts of the *TAW1*-CNS region (CR3, 24, and 25) exhibited severe panicle branching phenotypes. Deletion of all or large parts of the H1 region showed no significant effect on panicle branching (CR9, 10, 15, and 16). Conversely, deletion of all or part of the H2 region yielded a moderate increase in panicle branching (CR18 to 23). These results suggest that alteration of the H2 region is essential for phenotypic effects, whereas alteration of the H1 region enhances the effects of mutations in the H2 region. Notably, we found two independent CArG-like sequences (binding elements of MADS-box transcription factors) ^11^ embedded in H1 and H2, suggesting MADS transcription factor-mediated repression of *TAW1* transcription (Supplementary Fig. 7).

Although *taw1*-*D1*/-*D2* alleles showed substantial increases in spikelet number per panicle, they also decreased fertility ^8^, suggesting the need for a novel allele with moderately increased spikelet number per panicle as the ‘optimal’ allele. The genome-edited line with deletion of a part of the H2 region (CR19), displaying moderate increase in panicle branching, showed normal fertility while the line with large deletion in all part of the *TAW1*-CNS region (CR3) showed decreased fertility (Supplementary Fig. 8), implying that the CR19 is a candidate for the optimal allele. *TAW1*-CNS is conserved in many grass species; our results indicate that its precise genome editing could be useful for fine-tuning panicle branching to create the optimal allele in grass species. Yoshida *et al*. ^8^ reported that increased panicle branching in *taw1-D1/D2* is associated with transcriptional upregulation of *TAW1*; thus, the degrees of panicle branching phenotypes in the genome-edited lines in this study likely result from varying degrees of *TAW1* transcriptional upregulation. Natural variation (*locule number*; *lc*) or genome editing of a putative CArG element downstream of the *SlWUS* gene in tomato causes a dominant mutation with a subtle phenotype (increased locule number), although no transcriptional changes are detectable ^3^. In contrast, our study demonstrated marked transcriptional upregulation by deletions in the downstream region of the target gene through genome editing (Fig. 2). Similar to *SlWUS*, putative CArG elements in H1 and H2 of *TAW1*-CNS likely function as silencer elements (Supplementary Fig. 7). Notably, even though we identified two candidate CArG-like elements, our results suggest that different patterns of genome editing in, across, or proximal to these elements could produce different degree of phenotypes. This suggests that CNSs may be comprised of multiple *cis*-regulatory elements and their inter-elemental sequences, both of which may play important roles for fine-tuning transcription of proximal genes. Genome-wide identification and utilization of silencer modules would enable upregulation of proximal genes and broaden the range of genome editing applications. The molecular mechanism of transcriptional suppression by *TAW1*-CNS remains undetermined. Further investigation into whether the *TAW1*-CNS region is involved in negative transcriptional regulation via formation of intragenic chromatin loops, as previously reported ^12^, would be of interest. Consequently, CNSs around agronomically important genes could serve as useful target sites for genome editing to produce allelic series with variously altered gene expression and phenotypes to obtain ideal breeding materials.

## Methods

### Phylogenetic analysis

Predicted amino acid sequences and genomic sequences of *TAW1* in rice (*Oryza sativa* ssp. *japonica*) and its homologues in *Leersia perrieri, Brachypodium distachyon, Aegilops tauschii, Hordeum vulgare, Sorghum bicolor, Zea mays, Setaria italica, Panicum hallii* FIL2, *Musa acuminata, Dioscorea alata, Arabidopsis thaliana*, and *Marchantia polymorpha* were obtained from Ensembl Plants release 51 (https://plants.ensembl.org). *TAW1* and its homologues with their annotation IDs are listed in Supplementary Table 1. Amino acid sequences were aligned using CLC Workbench (v. 24.0.1) (Qiagen). Unrooted phylogenetic trees were generated by the neighbour-joining method with 1000 bootstrap replicates in CLC Workbench.

Phylogenetic shadowing of the *TAW1* regions was performed using mVISTA 13 (http://genome.lbl.gov/vista) in AVID mode. The analysis utilized regions from 1000 bp upstream of the predicted translational start site to 1500 bp downstream of the predicted translational stop codon.

### Identification of CNS

Chromosomes with *TAW1* orthologues from 9 Poaceae species were analysed: *O. sativa japonica* chromosome 10; *L. perrieri* chromosome 10; *B. distachyon* chromosome 3; *A. tauschii* chromosome 1D; *H. vulgare* chromosome 1H; *P. hallii*FIL2 chromosome 9; *S. italica* chromosome IX; *S. bicolor* chromosome 1; and *Z. mays* chromosome 5. Their soft-masked DNA sequences were retrieved from Ensembl Plants release 49 (https://plants.ensembl.org/).

Multiple alignment was performed mostly in accordance with the UCSC Genome Browser’s pipeline ^14^. First, chromosome-wide pairwise alignment of each species against the rice sequence was performed using LASTZ aligner ^15^ with the following parameters: --gap = “400,30” --xdrop = 910 --hspthresh = 3000 --ydrop = 9400 --gappedthresh = 3000 --inner = 2000. Local alignments were then extended and joined with axtChain (-minScore = 3000 -linearGap = “medium”). The best and nonoverlapping alignments were extracted with chainNet (-minSpace = 25 -minScore = 2000) and netToAxt. These pairwise alignments were integrated into multiple alignment blocks on the rice chromosome by MULTIZ ^16^ with the parameters R = 30 M = 18.

Sequence conservation was evaluated with PHAST tools ^17^. The nonconserved phylogenetic model was estimated from the fourfold degenerate (4d) sites using phyloFit. The gene annotation file from Ensembl Plants was used to identify 4d sites. Base-by-base conservation scores and discrete conserved elements were predicted after estimation of the scaling parameter ρ for the conserved model with phastCons (--target-coverage = 0.25 --expected-length = 12 -- most-conserved). CNSs were defined as conserved elements of at least 15 bp after subtracting coding sequences.. JBrowse 2 ^18^ was used to visualize the CNSs along with other genomic features.

### Plant material and growth conditions

*O. sativa* L. cv. Nipponbare was used to produce genome-edited rice. The previously reported *taw1*-*D2* mutant ^8^ was isolated from the nonautonomous DNA transposon *nDart1*-*0*–promoted mutant collection derived from the MK1 line. This line was selected in a cross between the *japonica* variety ‘Matsumoto-mochi’ and ‘Shiokari’ variegated *pyl* (*pale yellow leaf*) NIL carrying *nDart1*-*0* and an active autonomous transposable element ^19^. All rice seeds were surface-sterilized in 50% sodium hypochlorite solution with 0.01% Tween-20 for 30 min, rinsed with sterile water, placed on Petri dishes containing half-strength liquid Murashige and Skoog (MS) medium, and incubated in a growth chamber for at least 7 days. Seedlings were then transferred to soil in black vinyl pots (9 cm in diameter) until maturity. Plants grew in the greenhouse under LD conditions (14 h light at 28°C and 10 h dark at 25°C).

### Production of genome-edited rice

The vector pZNH2GTRU6 (manuscript in preparation) was used for genome editing via recognition of NGG as the PAM sequence. This vector, derived from plasmid pZ2028 ^20^, contains the hygromycin phosphotransferase (HPT) gene driven by the nopaline synthase promoter, *Streptococcus pyogenes* Cas9 driven by the modified rice polyubiquitin promoter, and guide RNAs (gRNAs) for editing target genes driven by the rice *U6-2* noncoding RNA promoter. The vector pZH_OsU6sgRNA_SpCas9–NGv1 ^21^ was used for genome editing via recognition of NG as the PAM sequence. One or two gRNAs were inserted into the vectors using the *Bbs*I site by ligation and/or In-Fusion HD Cloning (Takara). gRNA sequences are listed in Supplementary Table 2. Primers and oligonucleotides for construction are listed in Supplementary Table 3.

Transgenes were introduced into rice by *Agrobacterium tumefaciens*-mediated transformation as previously described ^22^. Hygromycin-resistant plants were transferred to soil in black vinyl pots (9 cm in diameter) until maturity. Genomic DNA was extracted from elongated leaves and subjected to PCR to confirm genome editing. Homozygous genome-edited plants were used for analyses, unless otherwise noted. Primers and oligonucleotides used to check genome editing are listed in Supplementary Table 3.

### RNA sequence analysis

Total RNA was extracted from immature inflorescence (3–6 mm in length) using an RNeasy Plant Mini Kit (Qiagen) and treated with DNase using a TURBO DNA-free Kit (Ambion) in accordance with the manufacturers’ protocols. RNA-seq libraries were prepared and sequenced on the Illumina NovaSeq 6000 platform at Novogene for paired-end 150-bp sequencing. Low-quality bases and adapter sequences in RNA-seq reads were removed by Trimmomatic v. 0.39 ^23^ with the following parameters: ILLUMINACLIP:TruSeq3–PE-2.fa:2:30:10 LEADING:15 TRAILING:15 SLIDINGWINDOW:10:15 MINLEN:50. Clean reads were mapped to the reference genome (Os-Nipponbare-Reference-IRGSP-1.0) by HISAT2 v. 2.2.1 ^24^ with the following parameters: --min-intronlen 20 --max-intronlen 10000. The latest RAP-DB (https://rapdb.dna.affrc.go.jp/) and RGAP (http://rice.uga.edu/) gene annotations as of March 11, 2022, were downloaded; the expression levels (transcripts per million; TPMs) of each gene were estimated using StringTie v. 2.2.1 ^25^. Differentially expressed genes (DEGs; FDR < 0.05) between WT and mutant lines were detected by DESeq2 ^26^.

### Prediction of transcription factor binding sites

Transcription factor binding sites in *TAW1*-CNS were predicted using the PlantRegMap database (https://plantregmap.gao-lab.org) ^27^. The *TAW1*-CNS sequence was entered into the Binding Site Prediction tool. Matched sequences with the lowest *P*-values (3.77 × 10^−6^ for TACCATAAAAAGCAA and 2.04 × 10^−5^ for TACCATAAGTAGCTA) in the input sequence were selected as top hit motifs.

## Data availability

Source data are provided with this paper. All additional data sets are available from the corresponding author upon reasonable request.

## Acknowledgements

We thank M. Kimizu, A. Nozaka, T. Akiyama, T. Kitashima, C. Isahaya, M. Nagakura, and M. Ishitsuka for assistance with production, genotyping, and phenotyping of genome-edited rice. We also thank M. Kuroda and M. Endo for providing the vectors Pznh2GTRU6 and Pzh_OsU6sgRNA_SpCas9–NGv1, respectively. This research was supported by a Research Program on Development of Innovative Technology grant (JPJ007097) from the Project of the Bio-oriented Technology Research Advancement Institution (BRAIN), JSPS KAKENHI Grant Number 22H00373, and by the Cross-ministerial Strategic Innovation Promotion Program (SIP) ‘Technologies for Smart Bio-industry and Agriculture’ (funding agency: BRAIN). This research was also partially supported by the Research Supporting Program of Research Center for Advanced Analysis, National Agriculture and Food Research Organization (NARO).

## Author information

## Contributions

T.K. and H.Y. designed the experiments and wrote the manuscript. T.K., F.L., and H.Y. analysed the phenotypic data. T.K., F.L., and S.C. performed the transgenic experiments and sequencing analyses. W.M.I. and T.M. performed analyses to detect CNSs. S.C. and Y.K performed RNA-seq analyses. A.Y. and J.K conceptualized the research. F.L., S.C., and J.K. edited the manuscript. All authors have reviewed and agreed to publication of the manuscript.

## Supplementary Figures

**Supplementary Fig. 1.**
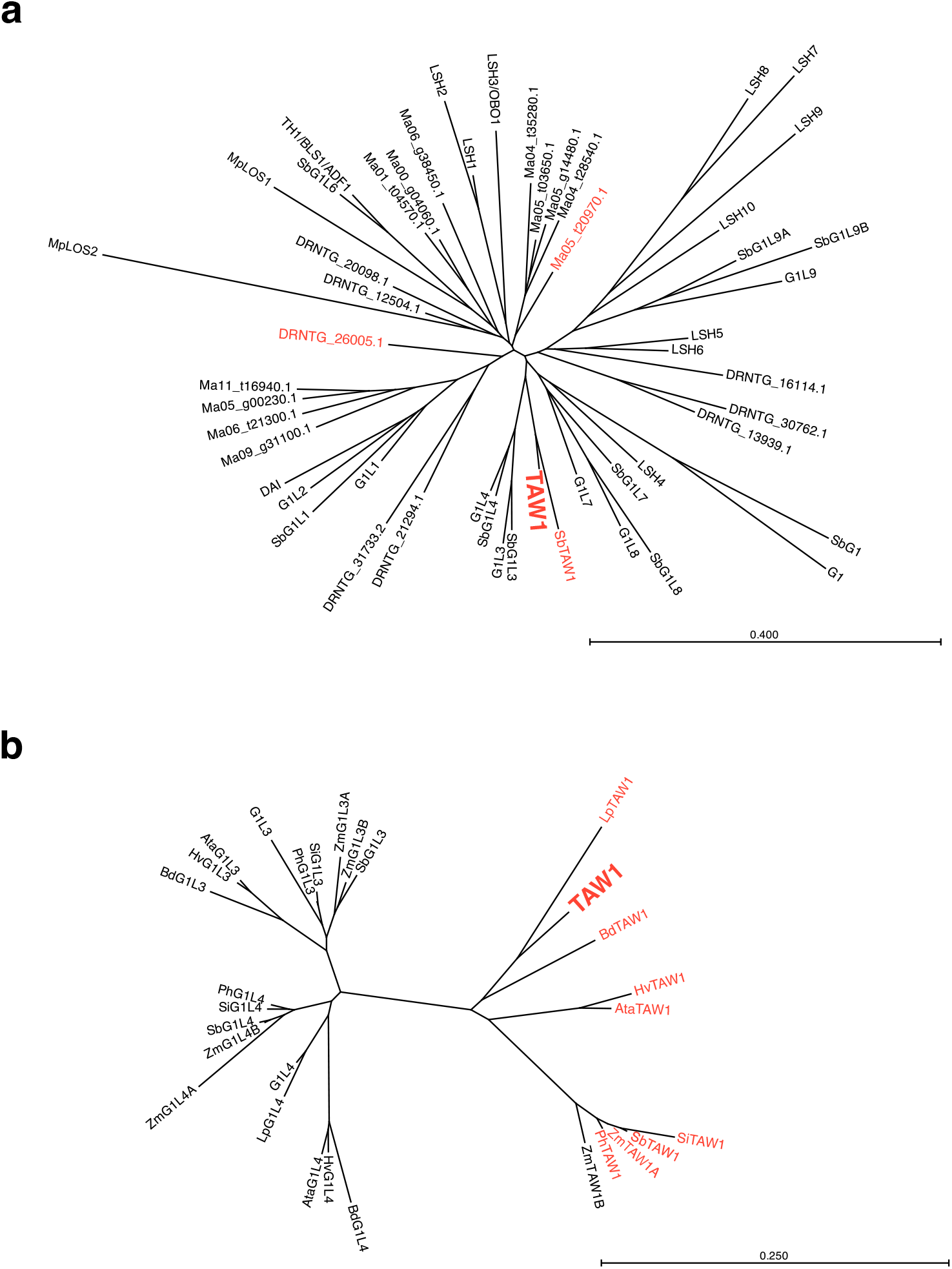
Phylogenetic tree based on predicted amino acid sequences of ALOG family genes. **a**, Phylogenetic tree of ALOG family proteins in rice, *Sorghum bicolor, Musa acuminata, Dioscorea alata, Arabidopsis thaliana*, and *Marchantia polymorpha*. **b**, Phylogenetic tree of TAW1, G1L3, and G1L4 proteins in 9 monocot species. Red symbols indicate proteins used for mVISTA and CNS analysis in Fig. 1a, b. Detailed information of proteins is listed in Supplementary Table 1. Numbers above branches indicate bootstrap values of 1000 trials (%) in **a** and **b**.

**Supplementary Fig. 2.**
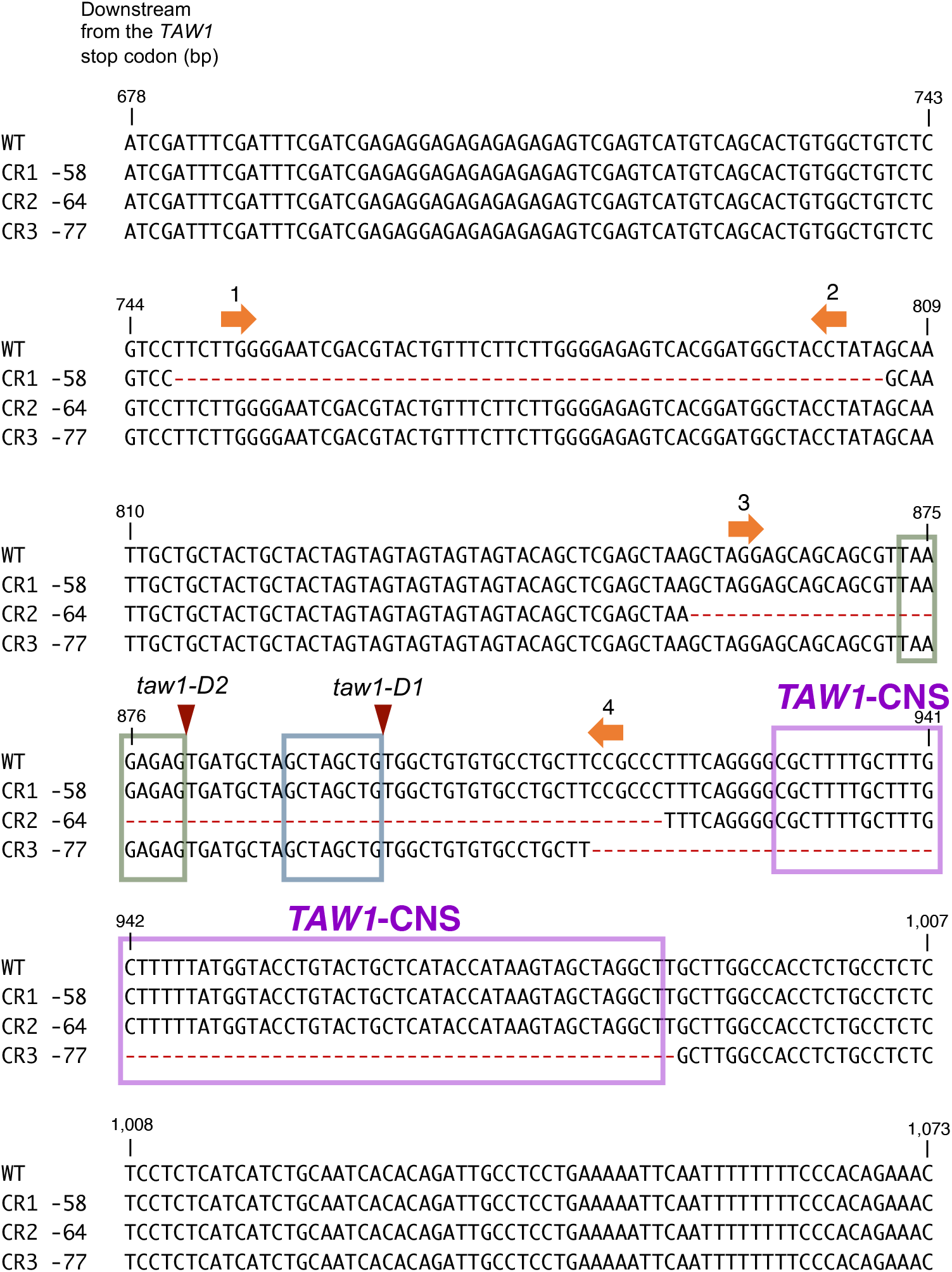
Positions of mutations, gRNA targets, and the CNS in the *TAW1* downstream region. The region 678–1073 bp downstream from the *TAW1* stop codon is shown. Positions of deletions in genome-edited lines (CR1 to 3) are shown as red dashes. Duplications in the *taw1-D1* and *taw1-D2* mutants are shown as green and blue boxes, respectively. Insertions in the *taw1-D1/D2* mutants are shown as red triangles. Positions of PAM sequences in gRNA targets (1 to 4) for genome editing are indicated by orange arrows.

**Supplementary Fig. 3.**
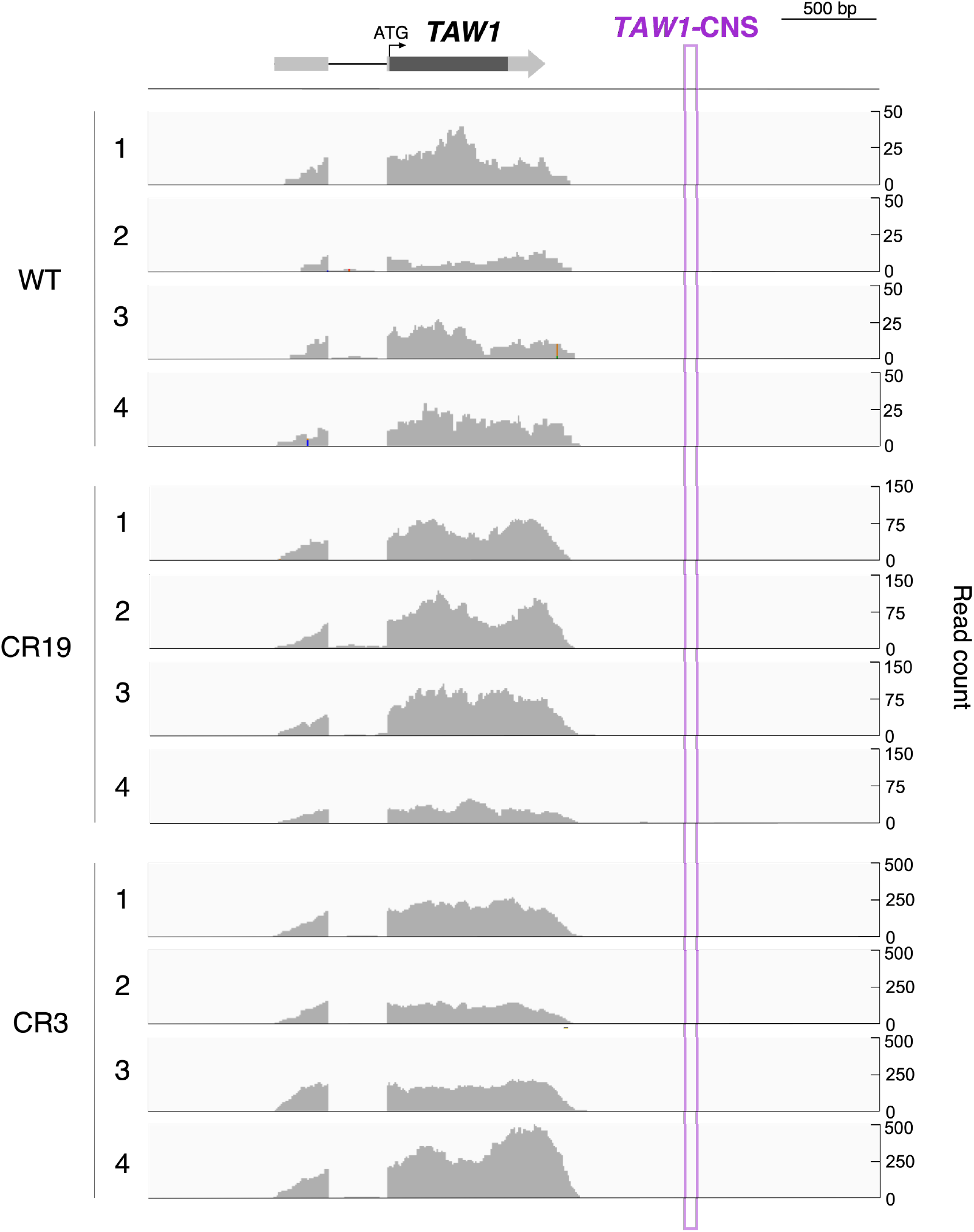
The *TAW1*-CNS region is not transcribed in young panicles of rice. Short reads were obtained from RNA-seq analysis in young panicle tissue of vector control (WT) and genome-edited lines (CR19 and CR3). Read counts from short-read mapping around the *TAW1* region were visualized by IGV browser (https://github.com/igvteam/igv). No reads mapped to the *TAW1*-CNS region in all samples. Results of 4 independent analyses of each line are shown. The annotated structure (RAP-DB) of *TAW1* is shown at the top.

**Supplementary Fig. 4.**
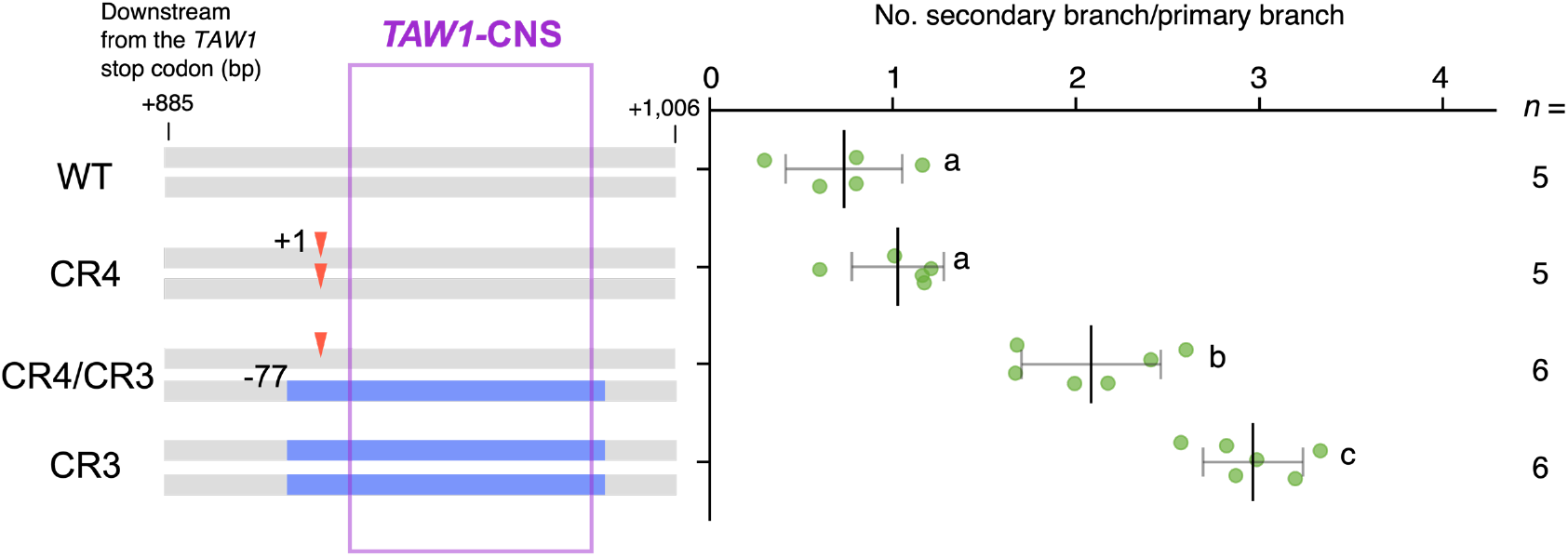
The panicle branching phenotype of *TAW1*-CNS deletion is inherited in a semidominant manner. Numbers of secondary branches per primary branch of a line transformed with vector control (WT), +1 homozygote (CR4), biallelic (CR4/CR3), and -77 homozygote (CR3) plants are shown. All plants were individuals segregated from the genome-edited line -77/+1 carrying the -77 and +1 biallelic mutations. Purple square indicates the *TAW1*-CNS region. Different letters in the right panels indicate statistically significant differences (*P* < 0.05; Tukey–Kramer test).

**Supplementary Fig. 5.**
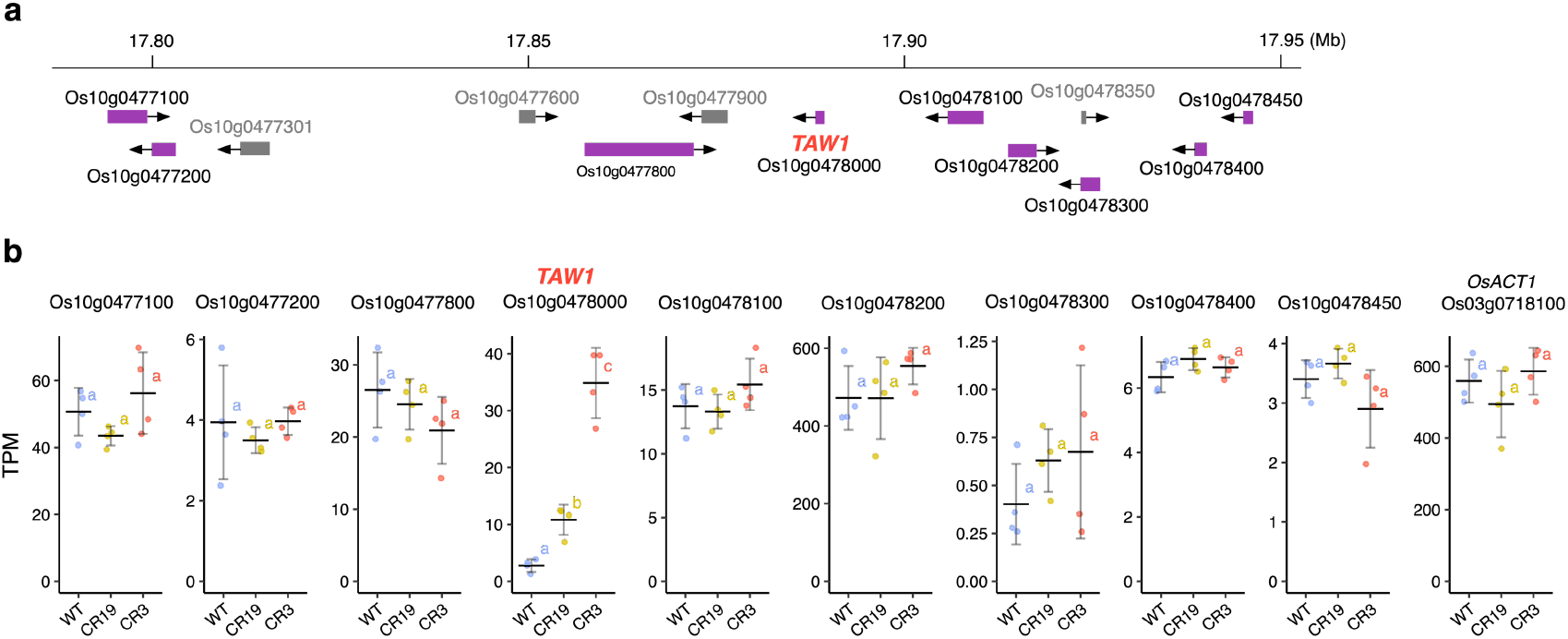
Transcription of *TAW1* and proximal genes in young panicles. **a**, Positions of *TAW1* and proximal genes in the rice genome. **b**, Normalized transcript expression levels (TPM) of *TAW1* and proximal genes in the immature inflorescence determined by RNA-seq analysis. Data of 9 genes (purple boxes in **a**) with TPM > 0.5 and *OsACT1* are shown. Data of Os10g0477800, *TAW1*, and Os10g0478100 are identical to Fig. 2d. Different letters indicate statistically significant differences (*P* < 0.05; Tukey–Kramer test) (*n* = 4).

**Supplementary Fig. 6.**
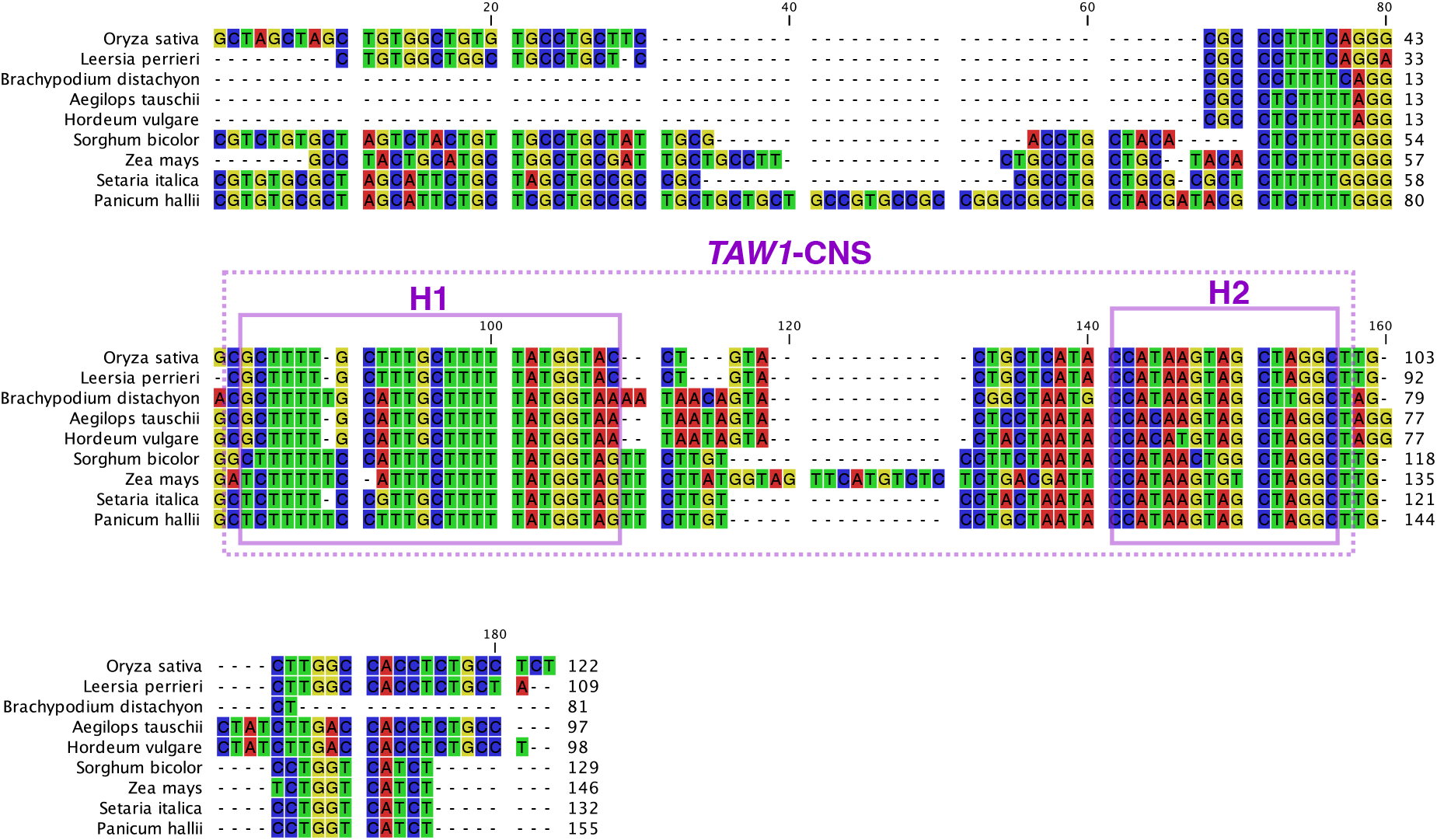
Genomic sequence alignment of *TAW1*-CNS region in 9 grass species. The region 885–1006 bp downstream from the *TAW1* stop codon in *O. sativa* was used to construct the alignment. Dashed box indicates *TAW1*-CNS detected in the BEP clade. Solid boxes indicate highly conserved CNS regions (H1 and H2) with conservation score ≥ 0.6 in the BEP clade.

**Supplementary Fig. 7.**
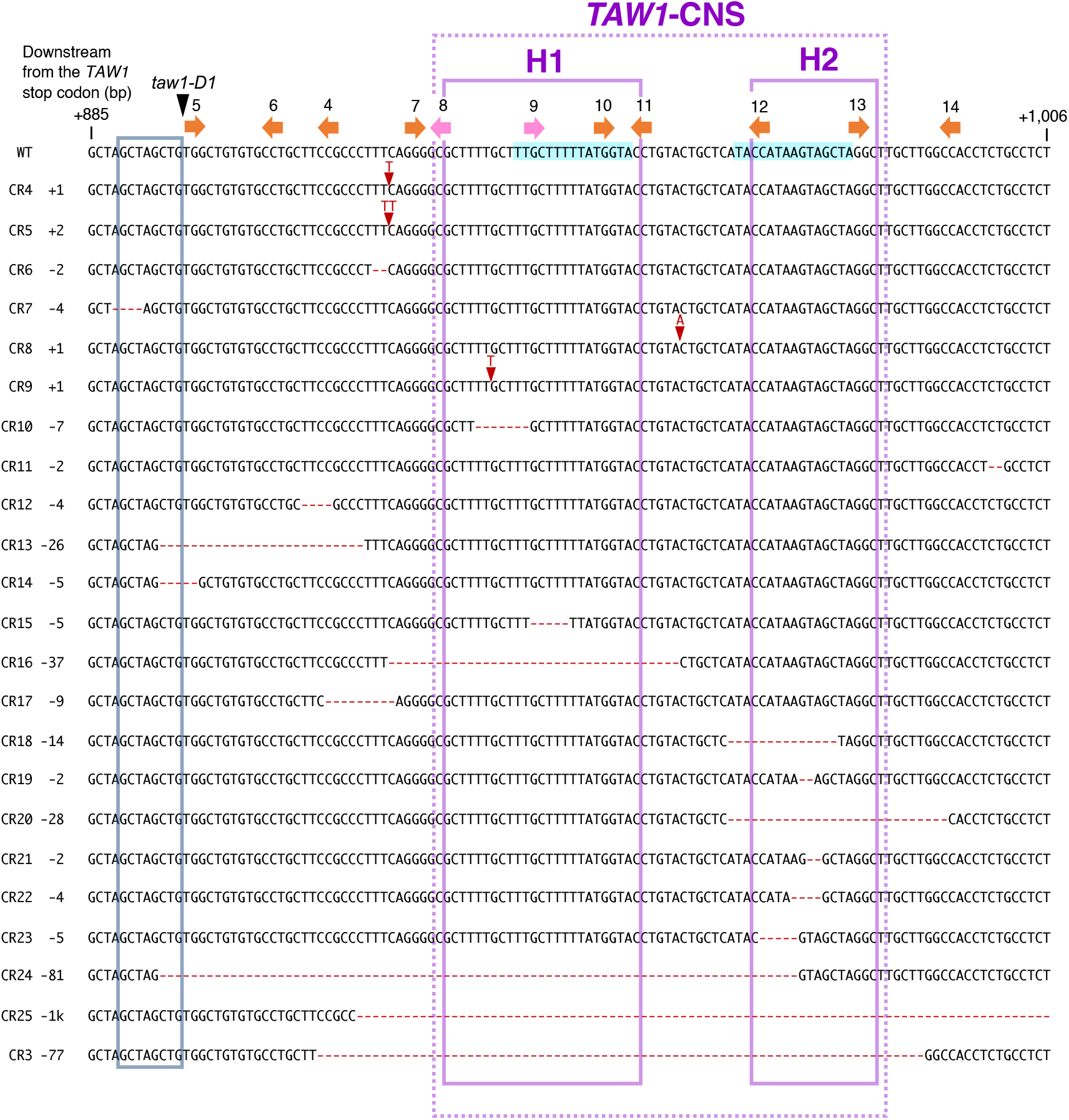
Positions of mutations and gRNA targets around the *TAW1*-CNS region. Positions of insertions or deletions in genome-edited lines (CR3 to 25) are indicated by red dashes and red triangles, respectively. Positions of PAM sequences in the gRNA target for genome editing, recognizing NGG or NG as the PAM sequence, are indicated by orange and pink arrows, respectively. A duplication and an insertion in the *taw1-D1* mutant are indicated by a light blue box and a black arrow, respectively. Light blue highlights in WT indicate CArG-box motifs predicted by PlantRegMap. The CR25 mutation (−1k) comprises a 1089-bp deletion and 5-bp (TTTTG) insertion.

**Supplementary Fig. 8.**
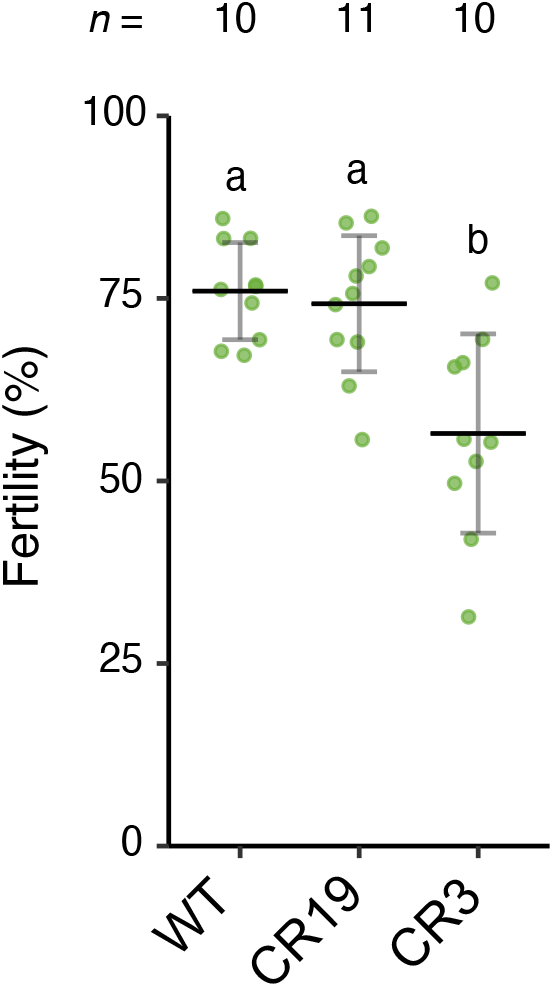
Fertility of the genome-edited lines in the *TAW1-*CNS region. Seed fertility rate in WT, genome-edited lines with moderate (CR19), and severe phenotype (CR3) for panicle branching were analysed. Different letters indicate statistically significant differences (*P* < 0.05; Tukey–Kramer test).

## Supplementary Tables

**Supplementary Table 1 *TAW1* and *TAW1* Homologues in Plants**.

**Supplementary Table 2 gRNA Sequences for Production of Genome-Edited Rice**.

**Supplementary Table 3 Primers and Oligonucleotides Used in This Study**.

